# Biological Factors and Statistical Limitations Prevent Detection of Most Noncanonical Proteins by Mass Spectrometry

**DOI:** 10.1101/2023.03.09.531963

**Authors:** Aaron Wacholder, Anne-Ruxandra Carvunis

## Abstract

Ribosome profiling experiments indicate pervasive translation of short open reading frames (ORFs) outside of annotated protein-coding genes. However, shotgun mass spectrometry experiments typically detect only a small fraction of the predicted protein products of this noncanonical translation. The rarity of detection could indicate that most predicted noncanonical proteins are rapidly degraded and not present in the cell; alternatively, it could reflect technical limitations. Here we leveraged recent advances in ribosome profiling and mass spectrometry to investigate the factors limiting detection of noncanonical proteins in yeast. We show that the low detection rate of noncanonical ORF products can largely be explained by small size and low translation levels and does not indicate that they are unstable or biologically insignificant. In particular, proteins encoded by evolutionarily young genes, including those with well-characterized biological roles, are too short and too lowly-expressed to be detected by shotgun mass spectrometry at current detection sensitivities. Additionally, we find that decoy biases can give misleading estimates of noncanonical protein false discovery rates, potentially leading to false detections. After accounting for these issues, we found strong evidence for four noncanonical proteins in mass spectrometry data, which were also supported by evolution and translation data. These results illustrate the power of mass spectrometry to validate unannotated genes predicted by ribosome profiling, but also its substantial limitations in finding many biologically relevant lowly-expressed proteins.

## Introduction

Ribosome profiling (ribo-seq) experiments indicate that genomes are pervasively translated outside of annotated coding sequences.^1^ This “noncanonical” translatome primarily consists of small open reading frames (ORFs), located on the UTRs of annotated protein-coding genes or on separate transcripts, that potentially encode thousands of small proteins missing from protein databases.^2^ Several previously unannotated translated ORFs identified by ribo-seq have been shown to encode microproteins that play important cellular roles.^3–6^ The number of translated noncanonical ORFs identified by ribo-seq analyses is typically very large, but many are weakly expressed, poorly conserved^7–9^, and not reproduced between studies^10^, suggesting that they may not all encode functional proteins. There has thus been considerable interest in proteomic detection of the predicted products of noncanonical ORFs.^11–15^ Detection of a noncanonical ORF product by mass spectrometry (MS) confirms that the ORF can generate a stable protein that is present in the cell at detectable concentrations and thus might be a good candidate for future characterization.

Over the past decade, numerous studies have attempted to identify noncanonical proteins using bottom-up “shotgun” proteomics in which MS/MS spectra from a digested protein sample are matched to predicted spectra from a protein database.^16,17^ These studies report hundreds of peptides encoded by noncanonical ORFs with evidence of detection in mass spectrometry data.^13–15,18–20^ However, these detections typically represent only a small fraction of the noncanonical ORFs found to be translated using ribo-seq. It is unclear whether most proteins translated from noncanonical ORFs are undetected by MS because they are absent from the cell, for example owing to rapid degradation, or because they are technically difficult to detect. Both the short sequence length and low abundance of noncanonical ORFs pose major challenges for detection in typical bottom-up MS analysis.^17^

Even given the low rates of detection of noncanonical proteins predicted by ribo-seq, there are suggestions^21,22^ that some of these claimed detections may be false positives, and true noncanonical detections even rarer. In particular, several practices in the statistical analysis of MS data might inflate the apparent number of confident noncanonical detections. Confidence is typically obtained in an MS analysis by controlling the false discovery rate (FDR, the expected proportion of inferred detected proteins that are incorrect). FDR estimation is usually estimated using a target-decoy approach, in which a set of proteins expected not to exist in the sample (“decoys”) are included in the sequence database along with predicted proteins (“targets”).^23^ As no decoys should be genuinely detected, the rate of inferred detection of decoys indicates the rate of false detections of targets. It is common to control FDR across the full proteome at 1%, such that the full list of detected proteins, including both canonical and noncanonical, contains only 1% false discoveries. However, this practice is recommended against by Nesvizhskii 2014^24^ and the Human Proteome Project.^25^ A strict FDR applied proteome-wide does not impose a strong constraint on the FDR among the noncanonical subset, and so the list of noncanonical detections may still contain a large proportion of false detections. This problem is exacerbated, moreover, if researchers control FDR at 1% on multiple datasets separately, and then merge the detected protein lists from each analysis. As true detections will tend to be shared between datasets while false detections will not, the FDR of the merged list is expected to be much higher than 1%.^26^ These problems can be addressed by setting a strict FDR on the noncanonical proteome specifically, and analyzing all datasets together in a single analysis. An additional potential problem comes from the manner in which decoy sets are constructed. Decoy sequences need to be unbiased, such that the MS analysis algorithm is just as likely to falsely claim a detection for a decoy or a target.^23,27^ Commonly used decoys constructed by reversing the sequence of target proteins have been shown to be unbiased for canonical proteins^28^, but it is unknown whether they are also unbiased for noncanonical proteins, and such a bias could cause incorrect estimation of FDR in either direction.

Several recent MS studies have aimed to improve detection of short, lowly-expressed proteins in *S. cerevisiae*. He et al. 2018^29^ used a combination of techniques to enrich for small proteins and detected 117 microproteins, including three translated from unannotated ORFs. Gao et al. 2021^30^ also used a combination of strategies to detect many small and low abundance proteins. Sun et al. 2022^31^ searched for unannotated microproteins in a variety of stress conditions and found 70, all expressed from alternative reading frames of canonical coding sequences. At the same time as these studies provided increased coverage of the yeast proteome, Wacholder et al. 2023^7^ integrated ribo-seq data from hundreds of experiments in over 40 published studies and assembled a high-confidence yeast reference translatome including 5372 canonical protein-coding genes and over 18,000 noncanonical ORFs. Here we leveraged these recent technical advances in MS and ribo-seq analysis to obtain a comprehensive, unbiased, account of noncanonical protein detection in *S. cerevisiae*, and investigate the biological and statistical factors affecting detection of noncanonical proteins.

## Results

### Noncanonical proteins and decoys detected at comparable rates

Using the MSFragger program^32^, we searched the three aforementioned published MS datasets optimized for detection of short, lowly expressed proteins^29–31^ against a sequence dataset that included all 5,968 canonical yeast proteins on Saccharomyces Genome Database (SGD)^33^ as well as predicted proteins from 18,947 noncanonical ORFs (including both unannotated ORFs and ORFs annotated as “dubious”) inferred to be translated in Wacholder et al. 2023^7^ on the basis of ribosome profiling data. The peptide-spectrum matches (PSMs) identified by MSFragger from all experiments among the three studies were pooled. FDR was estimated either for the full list of ORFs, or separately for canonical and noncanonical ORFs using a target-decoy approach.^23^ In both cases, we used the MSFragger expect scores, which indicates the confidence of the algorithm in each PSMs (with lower values indicating stronger matches), to estimate FDR at the protein level (number of decoy proteins passing threshold divided by number of target proteins passing threshold). A protein or decoy was considered detected if it had at least one unique PSM passing the threshold.

Among canonical ORFs considered alone, 4391 of 5968 had proteins detected at a 1% FDR (**Figure 1A**). For noncanonical ORFs considered alone, it was not possible to generate a substantial list of detected proteins at a 1% FDR because too many decoys were detected relative to targets at all confidence thresholds (**Figure 1B**). When the full proteome was considered together, 4389 proteins were found at a 1% FDR including 4371 canonical proteins and 18 noncanonical (**Figure 1C**). However, 10 noncanonical decoys also passed the 1% FDR expect score threshold, implying an estimated 56% FDR among these 18 noncanonical proteins. Thus, using a 1% proteome-wide FDR threshold, rather than a class-specific FDR strategy, results in a list of inferred noncanonical proteins of which a large fraction are false positives, as cautioned by <u>Nesvizhskii</u> 2014.^24^

**Figure 1:**
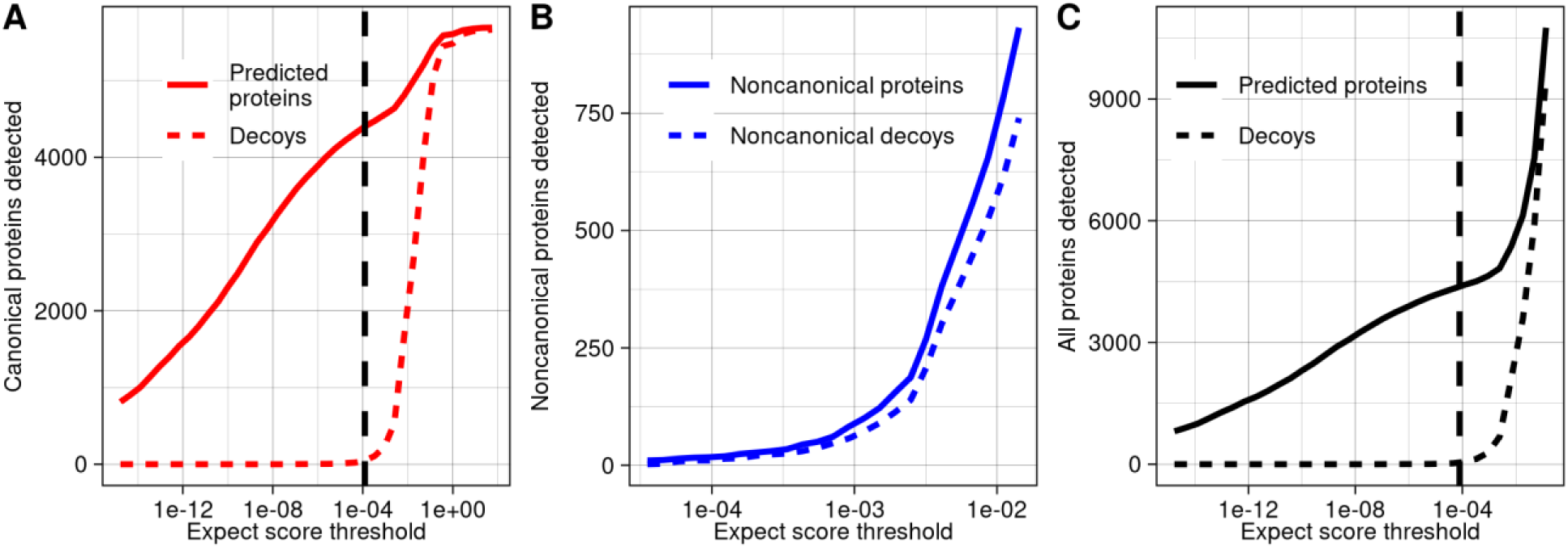
Few noncanonical proteins are confidently detected in MS data. A) The number of predicted proteins and decoys detected in MS data at a range of confidence thresholds among canonical yeast proteins. The dashed line signifies the 1% FDR threshold among canonical proteins. B) The number of predicted noncanonical proteins and decoys detected in MS data at a range of confidence thresholds. C) The number of predicted proteins and decoys detected in MS data at a range of confidence thresholds, considering noncanonical and canonical proteins together. The dashed line signifies the 1% proteome-wide FDR threshold. The data underlying this Figure can be found in S1 Data.

### Decoy bias among noncanonical ORF products leads to inaccurate FDR estimates

In general, there is a trade-off in target-decoy approaches such that setting a weaker confidence threshold results in a longer list of proteins inferred as detected, but with a higher FDR. In the case of yeast noncanonical proteins, the decoy/target ratio never went below 60% for any list of inferred detected target proteins larger than 10, and this ratio also did not converge to 1 even with thresholds set to allow 10,000 target proteins to pass (**Figure 2A**). The small enrichment of targets above decoys gives little confidence in detection of noncanonical ORF products at the level of individual proteins but leaves open the possibility that MS data could contain a weak biological signal.

**Figure 2:**
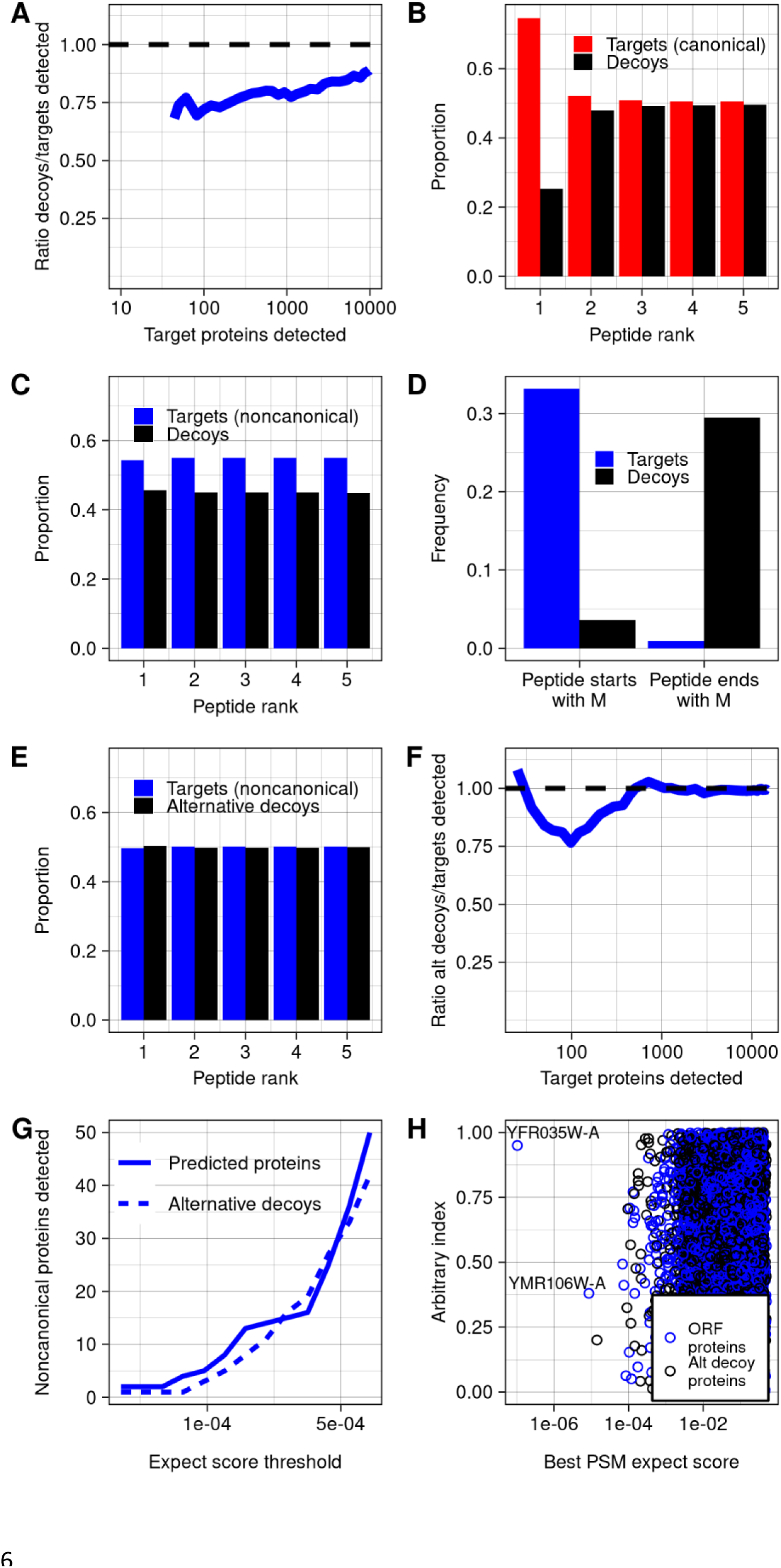
Decoy biases distort false discovery rate estimation. A) Among noncanonical proteins, the ratio of decoys detected to targets detected, across a range of targets detected, which varies with expect score threshold. Decoys are reverse sequences of the noncanonical protein database. B) Across all spectra, the proportion of peptide-spectrum matches of each rank that are canonical peptides vs. decoys. Peptide rank indicates the rank of the strength of the peptide-spectrum match, ordered across all peptides and decoys. C) Across all spectra, the proportion of peptide-spectrum matches of each rank that are noncanonical peptides vs. decoys. D) Among noncanonical ORF and decoy predicted trypsinized peptides that match spectra at any confidence level, the proportion that start or end with a methionine. E) Across all spectra, the proportion of peptide-spectrum matches of each rank that are noncanonical peptides vs. decoys, using the alternative decoy set. Alternative decoys are constructed by reversing noncanonical proteins after the starting methionine, such that all decoy and noncanonical proteins start with M. F) Among noncanonical proteins, the ratio of decoys detected to targets detected across counts of targets detected, using the alternative decoy set. G) The number of predicted proteins and decoys at a range of confidence thresholds, using the alternative decoy set. H) The best PSM expect scores for each noncanonical protein and decoy in the database, using the alternative decoy set. The data underlying this Figure can be found in S1 Data.

However, there is an alternative explanation for why targets are found at somewhat higher rates than decoys across a large range of confidence thresholds: decoy bias.^23^ The accuracy of FDR calculations require that target and decoy false positives are equally likely at any threshold, but this assumption could be violated if there are systematic differences between targets and decoys. Decoy bias has been assessed in previous work by comparing the number of target and decoy PSMs below the top rank for each spectra: if a peptide is genuinely detected, it will usually be the best match to its spectra, and so lower-ranked matched peptides will be false and should appear at approximately equal numbers for both targets and decoys.^23^ Among canonical ORFs, this expected pattern is observed (**Figure 2B**). In contrast, targets substantially outnumber decoys at all ranks for noncanonical ORFs (**Figure 2C**). We reasoned that this bias could be explained by the short length of noncanonical proteins. Indeed, many predicted peptides derived from noncanonical ORFs include the starting methionine, while decoys, consisting of reversed sequences from the protein database, are more likely to end with methionine (**Figure 2D**). To eliminate this large systematic difference, we constructed an alternative decoy database in which decoys for noncanonical proteins were reversed only after the leading methionine. When this database is used, the number of noncanonical targets and decoys at each rank is close to equal (**Figure 2E**) and the target/decoy ratio converges to one as confidence thresholds are lowered (**Figure 2F**). This behavior is consistent with expectations for a well-constructed decoy set. We therefore repeated our initial analysis using the alternative decoy set (**Figure 2G-H**) and used it for all subsequent analyses.

### Two noncanonical proteins show strong evidence of genuine detection

Using the alternative decoy set and standard MSFragger analysis, we remained unable to construct an FDR-controlled list of noncanonical proteins at a 10% FDR threshold because decoys were still detected at a similar rate as targets (**Figure 2G**). We therefore sought to examine the strongest hits to determine if we could identify evidence that any were genuine detections. Two noncanonical proteins had peptides with stronger expect scores than any decoys (**Figure 2H**; standard MSFragger approach in **Table 1; Supplementary Table 1**). We gave the ORFs encoding these proteins systematic names YMR106W-A and YFR035W-A following SGD conventions.^33^ A YFR035W-A peptides matched to two distinct spectra at thresholds stronger than the best decoy match (**Supplementary Figure 1**). Only a single YMR106W-A peptide was found at this threshold, but three additional YMR106W-A peptides had stronger matches than the next strongest decoy (**Supplementary Figure 2**). Moreover, YMR106W and YFR035W-A both had translation rates (in-frame ribo-seq reads per codon) greater than 99% of noncanonical ORFs in the Wacholder et al. 2023 dataset.^7^ The identification of multiple matching spectra for these noncanonical proteins and their relatively high rates of translation provide strong support that these are genuine detections. We note that this analysis also detected the three peptides from noncanonical ORFs reported by He et al. 2018^29^ with stronger expect scores than any decoys. However, as these proteins have recently been annotated by SGD as a result of the He et al. findings, they were not included in our noncanonical ORF set.

**Table 1:** Noncanonical ORFs possibly detected in mass spectrometry data.

| Systematic name | Approaches used to find | Coordinates | Peptides detected (spectra count) | Best MSFragger expect score | Quantile of translation rate | Evidence of conservation | Strength of evidence** |
| --- | --- | --- | --- | --- | --- | --- | --- |
| YMR106W-A* | Standard MSFragger, MS-GF+, MaxQuant | chrXIII:4809 24-481187 | ISMEAINNFIK (1) | 8.66e-06 | 0.9963 | None | Strong |
| YFR035W-A* | Standard MSFragger, MS-FG+ | chrVI:22626 0-226550 | HLNIPDLRFEK (2) | 1.04e-07 | 0.995 | Conserved within genus | Strong |
| YPR195C-A* | MaxQuant | chrXVI: 930265-930405 | LSAMVLTK (1) | 1.79e-04 | 0.61 | None | Weak |
| YOR109W-A* | Acetylation | ChrXV:<br>528606-528701 | LSLKC GCCSVCIPLNERR<br>(1) | 8.37e-06 | 0.27 | None | Weak |
| YIL059C | Non-enzymatic<br>end | chrIX:24655<br>0-246915 | EFD F D V G Y E E F V R (1) | 2.78e-06 | 0.88 | Conserved<br>with *S. jurei* | Strong |
| YNL155C-A* | Same-<br>strand<br>overlap | chrXIV:<br>341911-342135 | KQHTEWPIEENR (1),<br>MIGLIVVPILFAIK (11) | 5.77e-09 | 0.9963 | Conserved<br>within genus | Strong |

\*Assigned in this study.
\*\*Assessed based on proteomic, translation and evolutionary evidence

YMR106W-A is located 27 nt away from a Ty1 long terminal repeat. No homologs outside *S. cerevisiae* were found using BLASTP or TBLASTN against the NCBI non-redundant and nucleotide databases or against the 332 budding yeast genomes collected by Shen et al. 2018.^34^ It is thus plausible that this ORF was brought into the *S. cerevisiae* genome through horizontal transfer mediated by Ty1 retrotransposition.^35^ This is a similar origin to that of ERVK3-1, a human microprotein derived from an endogenous retrovirus.^36^ YFR035W-A overlaps the canonical ORF YFR035C on the opposite strand. Several known microproteins are expressed on the opposite strand of other genes^37,38^, so it is possible that both YFR035W-A and YFR035C are protein-coding genes. However, YFR035C was not detected in our canonical protein MS analysis. YFR035C deletion was reported to increase sensitivity to alpha-synuclein^39^, but this observation stemmed from a full ORF deletion that would also have disturbed YFR035W-A. While YFR035C has 2.5 in-frame ribo-seq reads per codon mapping to the ORF in the Wacholder et al. 2023^7^ dataset, YFR035W-A has 232, greater by a factor of 93 (**Figure 3A**). In a multiple sequence alignment with other species in the Saccharomyces genus, the full span of the YFR035W-A amino acid sequence aligns between all species (**Figure 3B)**, while other species have an early stop preventing alignment with most of the YFR035C amino acid sequence (**Figure 3C**). Thus, evolutionary, translation and proteomics evidence all indicate that unannotated ORF YFR035W-A is a better candidate for a conserved protein-coding gene than annotated ORF YFR035C.

**Figure 3:**
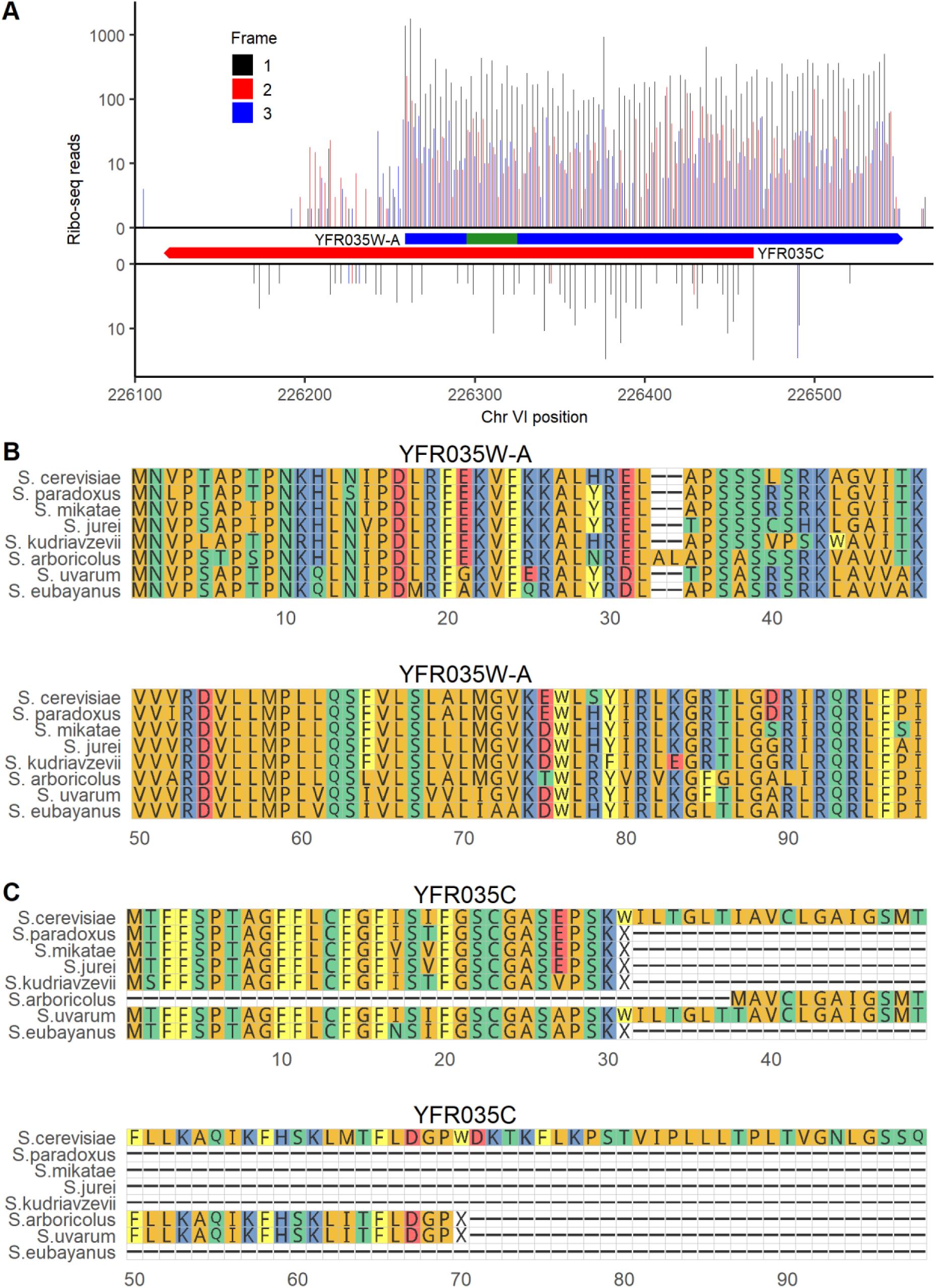
Translation and evolutionary evidence indicates that unannotated ORF YFR035W-A is likely a conserved gene. A) ribo-seq reads on unannotated ORF YFR035W-A (top) and annotated ORF YFR035C (bottom). The bounds of each ORF are indicated in boxes. The location of the detected peptide is indicated in green. Reads are assigned to the reading frame in which the position they map to is the first position in a codon; on each strand frame 1 corresponds to the reading frame of the ORF shown. The data underlying this Figure can be found in S1 Data. B) Alignment of the amino acid sequence of YFR035W-A with its homologs across the *Saccharomyces* genus. C) Amino acid alignment of the annotated ORF YFR035C and its homologs in *Saccharomyces*.

### Alternative strategies for MS search yield two additional noncanonical peptide detections

Aside from YMR106W-A and YFR035W-A, the standard MSFragger approach did not confidently detect proteins encoded by noncanonical ORFs supported by ribo-seq. We therefore considered some reasons we could miss noncanonical proteins present in the data and employed alternative approaches to test these possibilities. For each approach, we determined whether a substantial list of noncanonical ORFs could be constructed with FDR of 10% at the protein level. If not, we further investigated peptides with MSFragger expect scores < 10^-5^, similar to the level at which YMR106W-A was detected, or else the strongest candidates if another program was used.

First, we hypothesized that a mismatch between the environmental conditions in which the ribo-seq and MS datasets were constructed may explain the low number of detected noncanonical proteins. To investigate this possibility, we reduced our analysis to consider only ribo-seq and MS experiments conducted on cells grown in YPD at 30° C. The target/decoy ratio looked similar to the analysis on the full dataset, with no noncanonical protein detection list generatable with a 10% FDR (**Figure 4A**). The only noncanonical proteins detected at a 10^-5^ expect score threshold were the same two as in the standard analysis.

**Figure 4:**
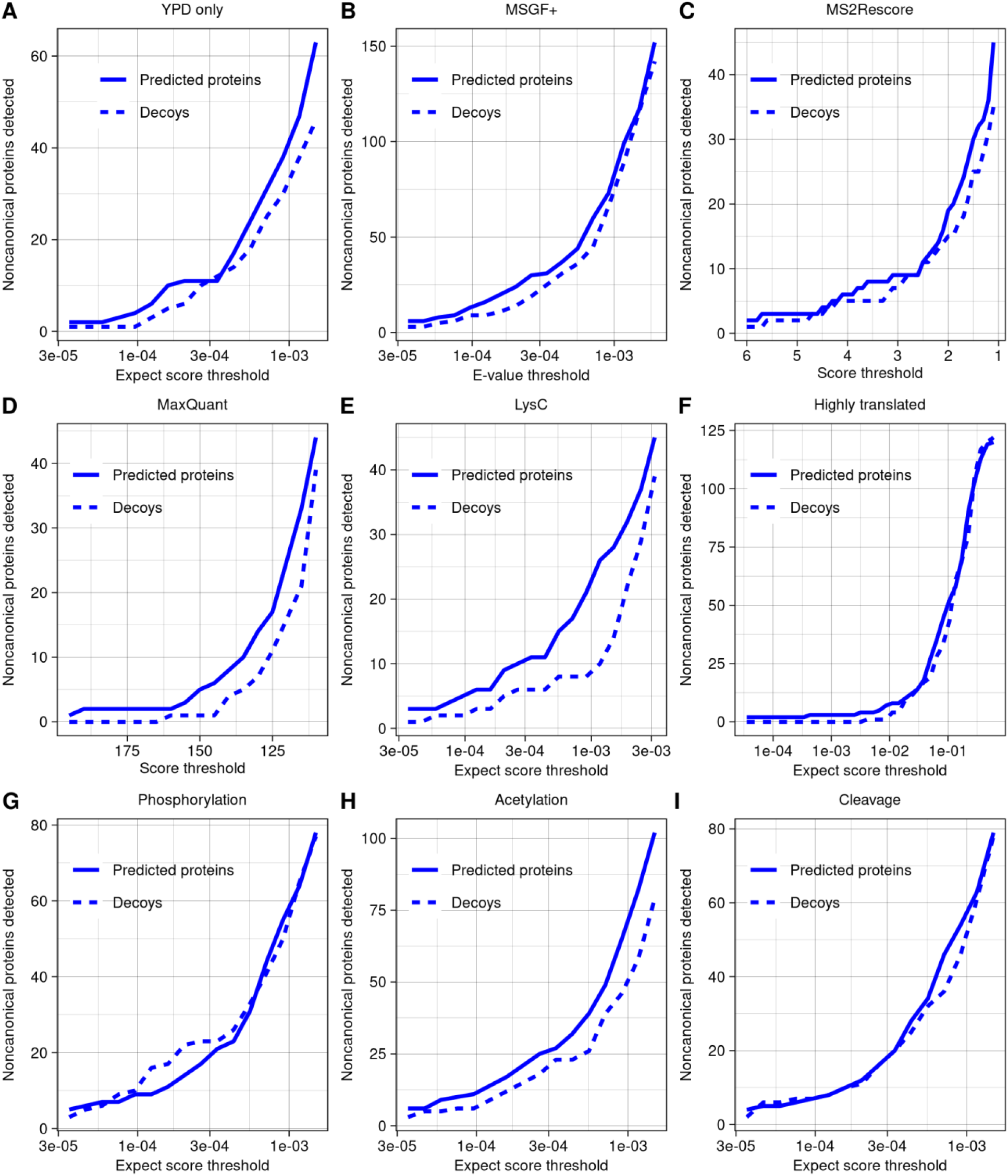
Alternative strategies for detecting noncanonical ORF products yield few additional discoveries. A-I) The number of predicted proteins and decoys detected across a range of thresholds, using a variety of strategies for detection. Aside from the specific changes indicated, all searches were run using the same parameter settings. The data underlying this Figure can be found in S1 Data. A) Analysis using only ribo-seq and MS data taken from yeast grown in YPD at 30° C. B) Analysis using the program MSGF+. C) Analysis using the rescoring algorithm MS^2^Rescore on MSGF+ results. Higher scores indicate higher confidence. D) Analysis using the program MaxQuant. Higher scores indicate higher confidence. E) Analysis including only experiments using LysC as protease. F) Analysis restricted to database of 379 predicted noncanonical proteins encoded by ORFs in the top 2% of in-frame ribo-seq reads per codon. G) Analysis allowing for phosphorylation of threonine, serine or tyrosine as variable modifications. H) Analysis allowing for acetylation of lysine or n-terminal acetylation as variable modifications. I) Analysis allowing detection of peptides with one end as a non-enzymatic cut site.

Next, to ensure our results were not specific to the search program MSFragger, we repeated our analysis using MS-GF+.^40^ The pattern of target vs. decoy detection was again similar to the standard MSFragger analysis, with no noncanonical detection list generatable with a 10% FDR (**Figure 4B**). The only noncanonical proteins detected at a 10^-5^ e-value threshold (e-value is the PSM confidence score given by MS-GF+) were YMR106W-A and YFR035W-A, also found by MSFragger. We then applied the machine learning based MS^2^Rescore algorithm^41^ to rescore the MSGF+ results, as this has been shown to improve peptide identification rates in some contexts. However, this also did not improve target-decoy ratios (**Figure 4C**). We also performed a search using MaxQuant^42^, which uses the Andromeda score^43^ to indicate the strength of a peptide-spectra match. The general pattern was similar to MSFragger and MS-GF+ (**Figure 4D**), with only three peptides given stronger scores than the strongest decoy; two belonged to YMR106W-A and one to a different hypothetical protein we named YPR195C-A following SGD conventions. However, this hypothetical protein was identified from a peptide found only once, showed no evidence of conservation in the *Saccharomyces* genus, and was not translated at high levels (**Table 1**); we therefore conclude that it may not be a genuine detection.

Work in other species has shown that use of multiple proteases, rather than trypsin alone, can improve detection of small or noncanonical proteins.^44,45^ We therefore investigated whether use of alternative protease could help with noncanonical detection in the dataset we examined. Some experiments in Gao et al. 2021^30^ used LysC as the enzyme, and though these were included in all analyses, all detections noted so far were tryptic peptides. When the LysC experiments were analyzed alone using MSFragger, we were still unable to construct a list of noncanonical proteins at 10% FDR (**Figure 4E**) and there were no PSMs with expect scores below 10^-5^.

One challenge in MS proteogenomics is that expanding searches to larger sequence database sizes raises the threshold for detection, which could limit discoveries.^46^ To reduce this challenge, we constructed a sequence database consisting only of the proteins expressed from the top 2% of noncanonical ORFs by translation rate. With only 379 proteins, this database is much smaller than the canonical yeast proteome, yet still we did not observe an improvement in the decoy/target ratio or any additional detections at a 10^-5^ expect score (**Figure 4F**).

Next, we hypothesized that noncanonical proteins could have been missed from our searches due to post-translational modification or cleavage. Allowing for phosphorylation of threonine, serine, or tyrosine as variable modifications did not improve the decoy/target ratio or yield detection of any noncanonical phosphorylated peptides at a 10^-5^ expect score threshold (**Figure 4G**). Adding acetylation of lysine or N-terminal acetylation as variable modifications did not improve target/decoy ratios overall **(Figure 4H**), but a single hit with an expect score of 8.37 x 10^-6^ was found, which we named YOR109W-A following SGD convention. However, this hypothetical protein was identified from a peptide found only once, showed no evidence of conservation in the *Saccharomyces* genus, and was translated at lower levels than other noncanonical protein detections (**Table 1**); we therefore conclude that it may not be a genuine detection.

Allowing for peptides to have one end that is not an enzymatic cut site to search for potential cleavage products did not improve target/decoy ratios overall (**Figure 4I**), but a single additional noncanonical peptide was identified with a relatively strong expect score of 2.78 x 10^-6^ (**Supplementary Figure 3**). This peptide was from the ORF YIL059C, annotated as “dubious” on SGD, indicating that, in the view of SGD, the ORF is “unlikely to encode a functional protein.” YIL059C is in the 88^th^ percentile of translation rate and 99^th^ percentile of length among noncanonical ORFs, at 366 nt (**Table 1**). It overlaps on the opposite strand the ORF YIL060W, classified as “verified” on SGD. However, the references listed in support of YIL060W are all based on full deletion experiments which would disturb both ORFs and therefore do not distinguish between them.^47–49^ YIL060W may have been considered the more likely gene as its ORF is longer, at 435 nt. But as in the case of YFR035C and YFR035W-A discussed above, both ribo-seq and MS data provide more support for the noncanonical ORF than the canonical ORF on the opposite strand: YIL059C has 14 in-frame ribo-seq reads per codon compared to only 0.48 in-frame reads per codon for YIL060W (**Figure 5A**), and YIL060W was not detected in our MS analysis of canonical ORFs. Given that the YIL059C peptide had one non-enzymatic end, we tested whether it could be a signal peptide using the TargetP program.^50^ YIL059C has a predicted signal peptide cleavage site corresponding exactly to the detected peptide (**Figure 5B**), providing additional support that this is a genuine detection. Searching for homologs using TBLASTN, BLASTP and BLASTN in the NCBI databases and TBLASTN and BLASTN in *Saccharomyces* genus genomes at a 10^-4^ e-value threshold, YIL059C and YIL060W have detected DNA homologs only in *Saccharomyces* species *S. paradoxus*, *S. mikatae* and *S. jurei*. There was an intact protein alignment of YIL059C between *S. cerevisiae* and *S jurei* (**Figure 5C**) while YIL060W has no homologs that fully align in any species (**Figure 5D**). YJL059C is located adjacent, and on the opposite strand, to a Ty2 long terminal repeat. These observations are consistent with a transposon-mediated horizontal transfer of YIL059C prior to divergence between *S. cerevisiae* and *S. mikatae*, followed by loss in *S. paradoxus* and *S. mikatae* and preservation in *S. cerevisiae* and *S. jurei*. We do not rule out a role for YIL060W, but all considered evidence provides greater support for the biological significance of YIL059C.

**Figure 5:**
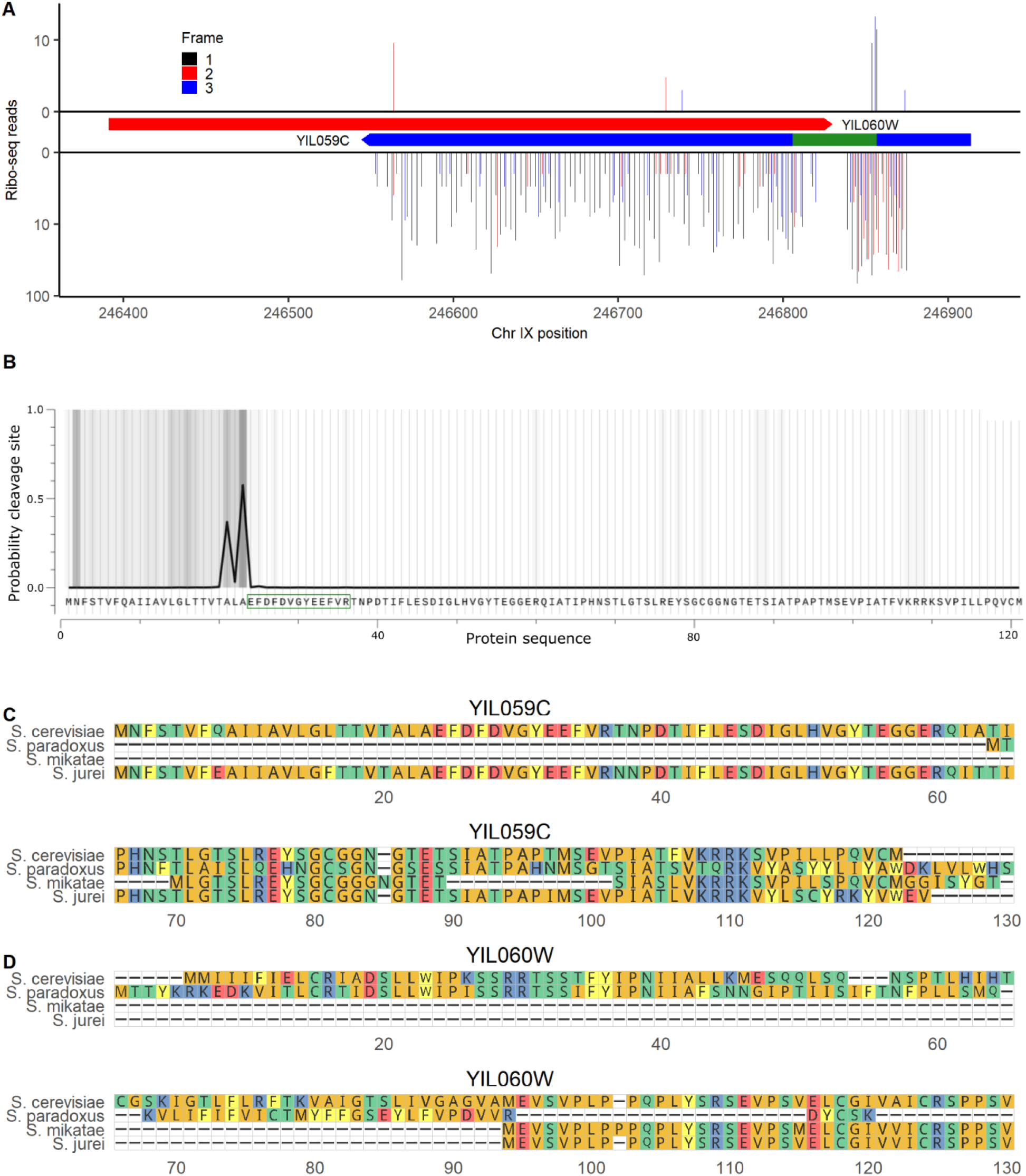
Dubious ORF YIL059C encodes a signal peptide. A) Ribo-seq reads on canonical ORF YIL060W (top) and “dubious” ORF YIL059C (bottom). The bounds of each ORF are indicated in boxes. The location of the detected peptide is indicated in green. Reads are assigned to the reading frame in which the position they map to is the first position in a codon; on each strand frame 1 corresponds to the reading frame of the ORF shown. B) Probability of a signal peptide cleavage site across the YIL059C sequence, as predicted by TargetP.^50^ The peptide detected in MS analysis is indicated by a green box. C) Alignment of YIL059C with the highest identity protein matches at the homologous locus in *Saccharomyces* species. Only species with a homologous locus (at the DNA level) are shown. D) Alignment of YIL060W, the canonical gene antisense to YIL059C, with its highest identity protein matches at the homologous locus in *Saccharomyces* species. The data underlying this Figure can be found in S1 Data.

Finally, we wanted to investigate a class of noncanonical ORFs not present in the Wacholder et al. translated ORF dataset: noncanonical ORFs that overlap a canonical ORF on the same strand. These ORFs are difficult to identify by ribo-seq because it is challenging to distinguish noncanonical ORF-associated ribo-seq reads from those of the canonical gene; however, some proteins encoded by noncanonical ORFs that overlap canonical ORFs have been identified in previous analyses^36,51^, including in the Sun et al. dataset included in our MS analysis.^31^ We therefore constructed a sequence database consisting of all canonical ORFs as well as noncanonical ORFs that overlap canonical ORFs on the same strand, with ORFs determined only from the genome sequence rather than expression evidence. Running this database against the full set of MS data, we again observed that, among noncanonical ORFs, decoys were detected at a high fraction of the rate of predicted peptides and so a list of confident noncanonical detections could not be established at reasonable false discovery rates (**Figure 6A).** These findings differ from those of Sun et al. 2022^31^, who found peptides from 70 noncanonical overlapping ORFs at a claimed 1% FDR. Of these claimed detections, 69 are also in our database, but none have peptides with stronger expect scores than the strongest decoys. To better understand this apparent discrepancy, we obtained the deposited MS program result output from the Sun et al. 2022 analysis. We observe that, within the Sun et al. results, the claimed noncanonical detections have confidence scores that are much weaker than canonical detections and similar to many decoys (**Supplementary Figure 4**). Thus, the Sun et al. results do not differ from ours because more high-confidence noncanonical PSMs were found. Rather, the difference is in statistical approach for FDR estimation. Sun et al. controlled FDR at a 1% proteome-wide level, rather than controlling a noncanonical-specific FDR as in our analysis. Moreover, Sun et al. analyzed multiple distinct datasets separately, each at a 1% FDR, and then constructed a combined list containing any protein found at 1% FDR in at least one analysis. Merging lists of detected proteins each constructed at 1% FDR is expected to generate a list with an FDR much higher than 1%.^26^

**Figure 6:**
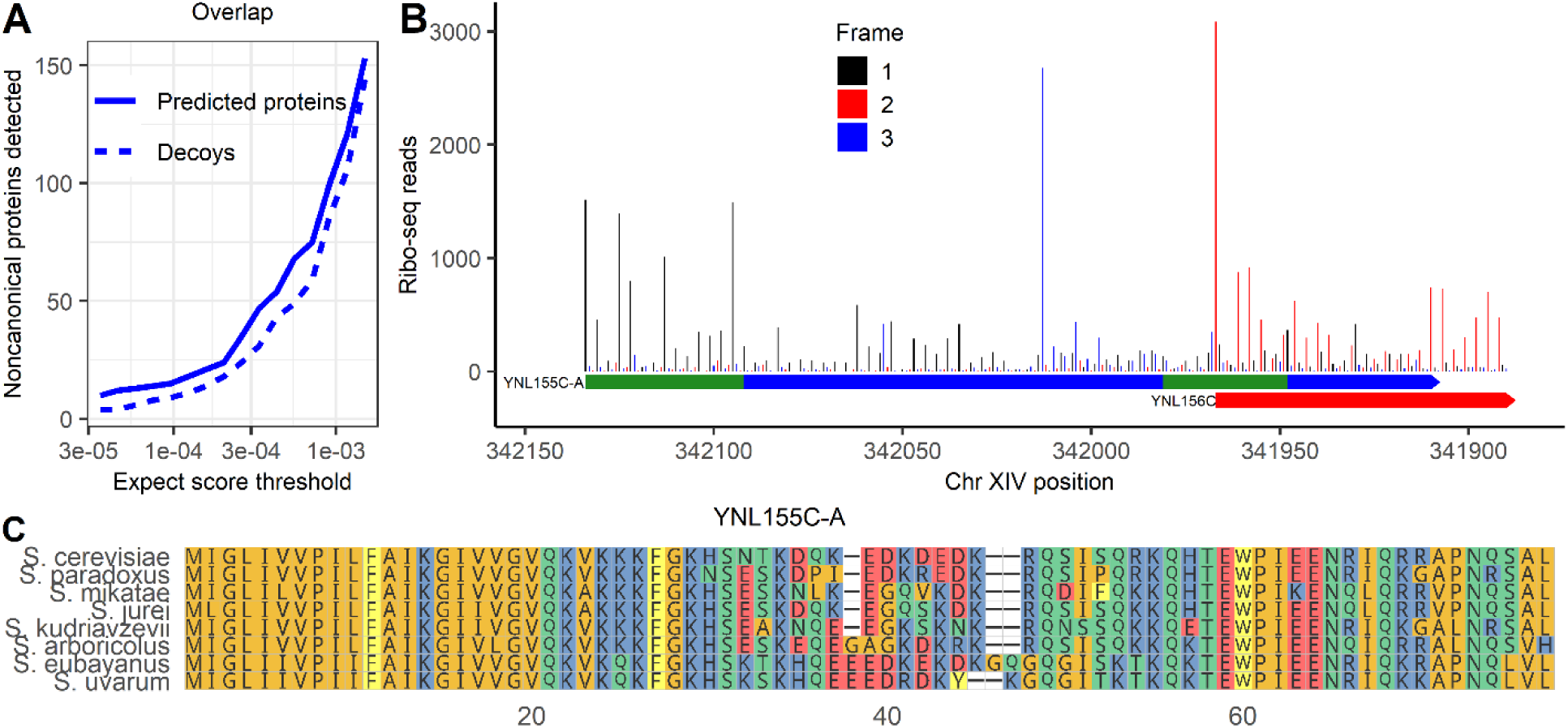
Noncanonical protein YNL155C-A, detected by MS, is well-translated and conserved in *Saccharomyces* genus. A) Predicted proteins and decoys detected in MS data at a range of expect-score thresholds, among noncanonical proteins that could be encoded by ORFs that overlap canonical ORFs on alternative frames. B) Ribo-seq reads across the YNL155C-A ORF. Reads are assigned to the reading frame in which the position they map to is the first position in a codon. Frame 1 is the reading frame of YNL155C-A. The full span of YNL155C-A and the start of YNL156C are shown. The position of the two peptides found in MS are in green. C) Multiple sequence alignment of YNL155C-A with its homologs in the *Saccharomyces* genus. The data underlying this Figure can be found in S1 Data.

In our analysis, only one overlapping ORF had associated PSMs with expect scores stronger than 10^-5^. We assigned it systematic name YNL155C-A following SGD conventions (**Table 1**).The stable translation product of YNL155C-A was supported by two distinct peptides which together were detected 12 times with expect scores below the best decoy score of 5.69 x 10^-7^, with the strongest value of 5.77 x 10^-9^ (**Supplementary Figure 5**).This 255 bp ORF overlaps canonical gene YNL156C for 57 of 255 bases. Its translation product was not identified in the Sun et al. analysis.^31^ A clear pattern of ribo-seq read triplet periodicity was observed in the frame of YNL155C-A (i.e., reads tend to match to the first position of a codon) before the overlap with YNL156C, indicating translation in this frame (**Figure 6B**). There also appears to be a triplet periodic pattern in a frame distinct from both YNL156C and YNL155C-A at the locus, suggesting that all three frames may be translated. Excluding the overlapping region, there are 265 reads per codon on the ORF that map to the first position of a codon in the YNL155C-A reading frame; this would put it in the 99.6^th^ percentile of translation rate among translated noncanonical ORFs in the Wacholder et al. dataset. No homologs were found in more distantly related species in a TBLASTN search against the NCBI non-redundant protein database, but YNL155C-A was well conserved across *Saccharomyces* (**Figure 6C**). As only 19 of 75 codons of YNL155C-A overlap YNL156C-A (**Figure 6B**), the strong amino acid conservation across the length of the full protein (**Figure 6C**) indicates purifying selection on YNL155C-A itself. Thus, proteomic, translation and evolutionary evidence all support YNL155C-A as a protein-coding gene.

### The low detectability of noncanonical proteins can be explained by their short lengths and low translation rates

We sought to understand why the large majority of proteins predicted from translated noncanonical ORFs remained undetected across multiple computational search strategies. A major difference between canonical and noncanonical proteins is length: the average canonical protein is 503 residues compared to only 31 among noncanonical proteins. Short size can affect protein detection probability through distinct mechanisms: shorter sequences provide fewer distinct peptides when digested, and the sample preparation steps of the MS experiment may be biased against small proteins.^17^ To investigate the first possibility, we computationally constructed all possible tryptic peptide sequences that could be theoretically detected from the proteins in the sequence database given their length and mass. Canonical proteins have an average of 62 theoretically detectable tryptic peptides compared to 6.7 for predicted noncanonical proteins. Among noncanonical proteins, 2,496 of 18,947 (13%) lack any theoretically detectable tryptic peptide, meaning these would be impossible to detect using our search strategy; by contrast only 11 canonical proteins (0.2%) lack any possible peptides. Not only are many noncanonical proteins undetectable due to the complete absence of potential tryptic peptides, but many others have so few potential peptides that it is unlikely that at least one will be discovered at current sensitivities. Indeed, the overall detection rate for canonical peptides is only 6% (at a 10^-6^ MSFragger expect score threshold). While 69% of canonical proteins have at least one peptide detected at this threshold, based on simulations, only 22% of noncanonical proteins would have a detectable peptide at this detection rate. These results illustrate how the short length of noncanonical proteins and the low numbers of potential tryptic peptides that result limit noncanonical detection.

Still, the larger challenge in noncanonical protein detection is not just the low number of possible peptides but the much lower detection rate among them than among canonical peptides. Our analyses only detect a handful of noncanonical proteins, far below the 22% that would be expected if lack of potential tryptic peptides was the only limitation. This is because, as a group, noncanonical proteins almost completely lack the high-confidence PSMs that support numerous canonical protein detections (**Supplementary Figure 6**).We therefore hypothesized that technical biases other than the number of potential tryptic peptides further limit the MS detectability of small proteins.

To investigate this hypothesis, we calculated the peptide detection rate, out of all theoretically detectable peptides, among different ORF size classes (**Figure 7A**). We observe a division between canonical ORFs shorter vs. longer than 150 nt. Among 27 canonical yeast ORFs shorter than 150 nt, none of 280 theoretically detectable peptides were detected at a 10^-6^ expect score threshold. This detection rate is significantly below expectation given the overall 6% rate at which canonical peptides are detected (binomial test, p = 4.73 x 10^-8^), suggesting that there may be technical biases limiting detection of proteins that are this short. As 83% of noncanonical ORFs (15,717) are shorter than 150 nt, short length can partially explain the low detectability of noncanonical ORF peptides. In contrast, however, among canonical ORFs longer than 150 nt, shorter lengths were associated with higher probabilities that a peptide was detected (**Figure 7A**). This is likely due to a trend of higher translation rates among shorter ORFs (**Supplementary Figure 7A**), which is also observed among noncanonical ORFs (**Supplementary Figure 7B**). This observation suggests that short size should not be a barrier to detection of peptides encoded by noncanonical ORFs longer than 150 nt. There are 3,080 such ORFs, potentially encoding 35,392 detectable peptides, yet only one peptide was found at a 10^-6^ expect score threshold (the peptide from YFR035W-A, **Table 1**).

**Figure 7:**
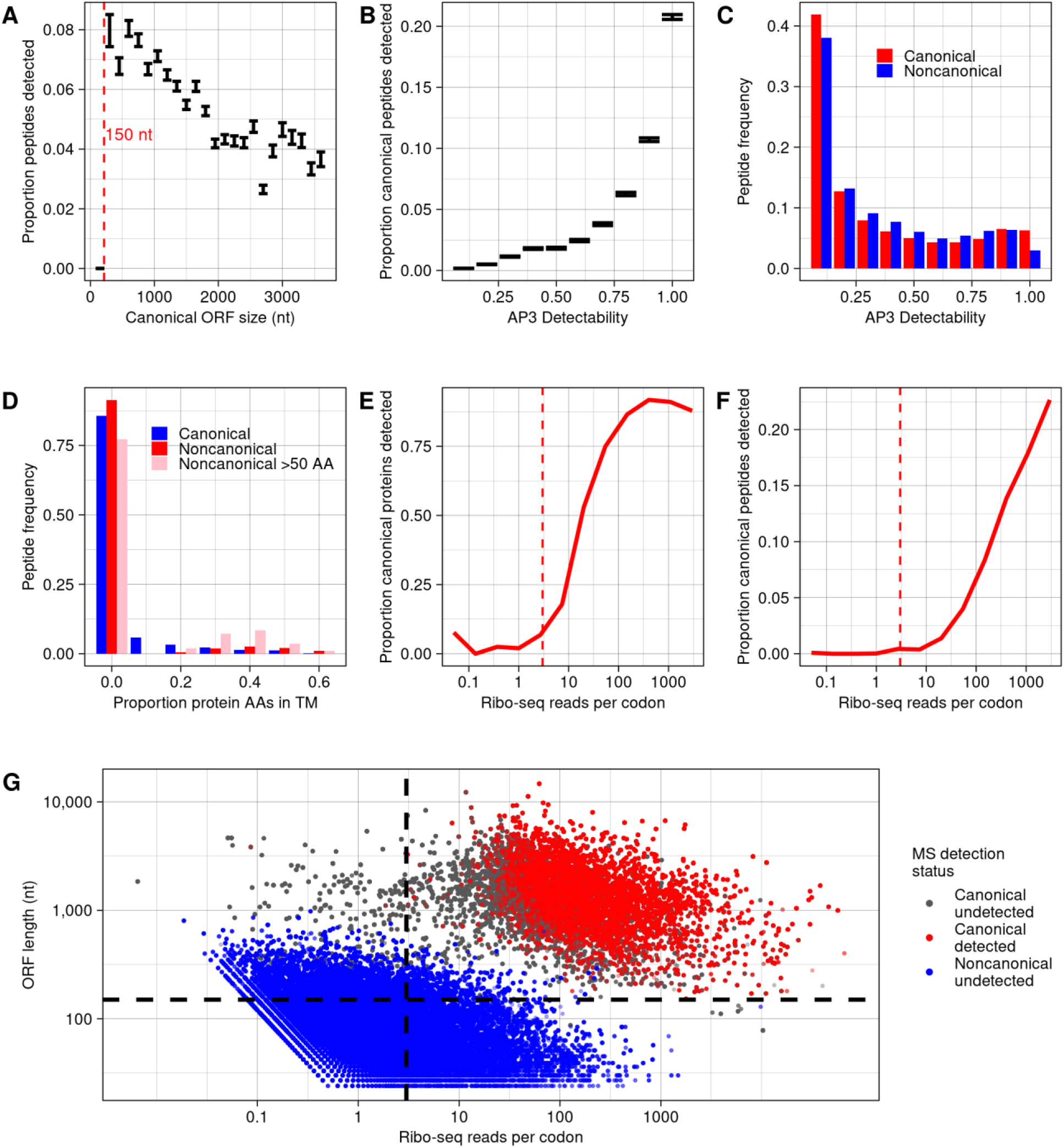
Lack of detection of noncanonical proteins can be largely explained by their low translation rate. A) The proportion of canonical peptides detected, among all eligible for detection, for ORFs of different size classes. Bars indicate a range of one standard error. A dashed line is drawn at 150 nt, below which no canonical peptides are detected. B) The proportion of peptides detected, among all eligible for detection, for canonical proteins binned by detectability score given by the AP3 algorithm.^53^ Bars indicate a range of one standard error. C) Frequencies of predicted peptides by detectability score among canonical and noncanonical proteins. D) Frequencies of predicted peptides among canonical proteins, noncanonical proteins, and noncanonical proteins larger than 50 amino acids, with proteins binned by proportion of amino acids in predicted transmembrane domains. Predictions were made using TMHMM.^54^ The first bin includes only proteins with no transmembrane domain predicted. E) Proportion of canonical proteins detected within bins defined by in-frame ribo-seq reads per codon mapping to the ORF. A dashed line is drawn at 3 reads per codon, below which few canonical proteins are detected. F) Proportion of canonical peptides detected, out of all eligible, within bins defined by in-frame ribo-seq reads per codon. A dashed line is drawn at 3 reads per codon, below which few canonical peptides are detected. G) For all peptides predicted from canonical and noncanonical translated ORFs with detectable mass and length, the in-frame ribo-seq reads per codon and ORF length is plotted. Each peptide is classed by whether it is canonical or noncanonical, and whether it is detected at a 10^-6^ expect score threshold. Nearly all detectable peptides are restricted to the top right section bound by dashed lines, where ORF length > 150 nt and reads per codon > 3. The data underlying this Figure can be found in S1 Data.

In addition to length, noncanonical ORFs also differ from canonical ORFs in amino acid composition.^7^ The amino acid composition of noncanonical proteins could limit detectability relative to canonical proteins because detectability of a peptide in an MS experiment is affected by its physical properties.^52^ To test this possibility, we applied the AP3 algorithm^53^, which predicts peptide detectability from peptide sequence, assigning a score from 0 to 1, to the full set of tryptic peptides predicted from the yeast translatome. As expected, detectability scores corresponded strongly to observed detection rates among canonical peptides (**Figure 7B**). For example, 20% of canonical peptides scoring above 0.9 were detected at a 10^-6^ expect score threshold, compared to only 0.17% of peptides scoring below 0.1, an 85-fold increase. The distribution of detectability scores was similar between canonical and noncanonical peptides overall, with the major difference being that 6.3% of canonical peptides scored above .9 compared to only 3% of noncanonical peptides (**Figure 7C**). If canonical peptides had the same distribution of scores as noncanonical peptides, the number of detected canonical peptides would be 83% of those found in actuality. Thus, amino acid composition does increase the difficulty in detection of noncanonical peptides, but this is a relatively small effect.

Transmembrane proteins are also more difficult to detect by MS.^55^ To determine whether a high transmembrane propensity among noncanonical proteins could help explain their low detection rates, we used TMHMM^54^ to predict transmembrane domains among all proteins. As expected, detectability of canonical peptides declines with increased transmembrane content of the protein (**Supplementary Figure 8**). Among noncanonical proteins overall, only 11% are predicted to have transmembrane domains, below the 20% of canonical proteins predicted to have one. However, among noncanonical proteins longer than 50 amino acids there is an excess of proteins with transmembrane domains that make up more than 20% of the protein (**Figure 7D**; p<10^-16^, chi-squared test). Thus, for some larger noncanonical proteins we would otherwise expect to be more likely to be detected, transmembrane domains likely hinder their detection.

Besides length and sequence composition, a major difference between canonical and noncanonical ORFs is expression level, and this too can affect the probability a protein is detected in MS data.^17^ We therefore evaluated the relation between translation level and detection probability using the ribo-seq data from Wacholder et al. The number of in-frame ribo-seq reads per codon that map to a canonical ORF is strongly associated with the probability of detecting the ORF product at a 10^-6^ expect score threshold, at both the protein (**Figure 7E**) and peptide (**Figure 7F**) level. As with protein length, we can use the canonical ORFs to infer an approximate detection limit: among 439 canonical ORFs with fewer than 3 in-frame reads per codon, only 3 of 9,253 theoretically detectable peptides were detected at a 10^-6^ threshold. Thus, almost all canonical peptides, with only these three exceptions, are found among ORFs with reads per codon above 3 and longer than 150 nt. Yet, only 448 noncanonical ORFs (2.4% of total) are in this category (**Figure 7G**). Thus, almost all noncanonical ORFs are outside the limits in which canonical ORF products are detected by MS.

For the 448 noncanonical translated ORFs displaying length and expression levels amenable to detection (longer than 150 nt and at least three reads per codon), we estimated the probability a peptide would be detected at a 10^-6^ expect score threshold under the assumption that detection probability depends only on translation rate. This probability was estimated as the peptide detection rate among canonical ORFs with a similar translation rate to the transient ORF (a natural log of reads per codon within 0.5).

Given these estimates, the expected total count of detected peptides for the 448 ORFs was 5.41. In reality, a single peptide was detected (the peptide from YFR035W-A, **Table 1**). To see whether observing only a single detection was surprising, we simulated the distribution of peptide detection counts under the estimated detection probabilities. The 95% confidence interval of noncanonical peptide detections ranged from 1-10. Thus, the single observed detection of a noncanonical peptide at a 10^-6^ expect score threshold is within range of expectations.

### Evolutionarily novel ORFs missed in MS data due to low sensitivity

Wacholder et al. identified a class of rapidly evolving, evolutionarily novel ORFs termed “transient ORFs.” Despite lacking long-term evolutionary conservation, transient ORFs can express proteins that have major effects on phenotype.^7^ Of 18,947 noncanonical ORFs analyzed here, 17,471 (91%) are inferred to be evolutionarily transient in the Wacholder et al. dataset; an additional 103 canonical ORFs are also classified as transient. As evolutionarily transient ORFs comprise such a large portion of the translatome, it is of interest to determine whether their products can be detected by shotgun mass spectrometry. No evolutionarily transient noncanonical ORF peptides were detected in our analyses, as none of the noncanonical proteins we identified (listed in Table 1) were classified as evolutionarily transient. Among the 103 evolutionarily transient canonical ORFs, none were detected at a 10^-5^ expect score threshold, and similar numbers of ORFs and decoys were found at weaker thresholds (**Supplementary Figure 9**).

Five transient canonical ORFs have been characterized in some depth^7^, including MDF1, a well-established *de novo* gene specific to *S. cerevisiae* that plays a role in the yeast mating pathway.^38^ Yet none of these show any evidence of detection in the MS datasets examined here, with expect scores far higher than what would constitute even weak evidence (**Table 2**). These results indicate that MS detection appears to miss the entire class of evolutionary transient ORFs, whether canonical or not, including even those known to play important biological roles.

**Table 2.**

| Canonical transient ORF | Major publication | Minimum expect score |
| --- | --- | --- |
| MDF1 | Li et al. 2010<sup>38</sup> | 1.85 |
| YBR196C-A | Vakirlis et al. 2020<sup>56</sup> | .99 |
| HUR1 | Omidi et al. 2018<sup>57</sup> | 1.62 |
| YPR096C | Hajikarimlou et al. 2020<sup>58</sup> | 0.10 |
| ICS3 | Alesso et al. 2015<sup>59</sup> | 0.03 |

## Discussion

Bottom-up mass spectrometry is an attractive approach for validating noncanonical ORFs supported by ribosome profiling due to the ease of testing large lists of predicted proteins but is limited by low sensitivity. Analyzing three mass spectrometry experiments optimized to find small proteins, we identified three noncanonical proteins expressed from ORFs identified as translated in a recent analysis of yeast ribosome profiling studies (YMR106W-A, YFR035W-A, and YIL059C). We additionally found MS evidence for an ORF not initially identified by ribo-seq, YNL155C-A, due to overlapping a canonical ORF on the same strand. All four proteins were translated at rates much higher than typical noncanonical ORFs, providing independent evidence that they are genuine protein-coding genes; three also showed evidence of evolutionary conservation. These findings illustrate the power of using proteomic, translation, and evolutionary evidence in combination to identify undiscovered genes at high confidence even in a well-annotated model organism.

Nevertheless, the vast majority of ribo-seq supported noncanonical ORFs showed no evidence of detection in MS datasets. We show that the low rates of detection of noncanonical ORFs can be explained primarily by their short size and low translation rate: canonical ORFs at similar sizes and translation rates are also very rarely detected. The general amino acid composition of noncanonical ORFs, and the abundance of transmembrane domains among the longest ones, further contribute to hindering detection. As these factors explain the differences in detectability between canonical and noncanonical ORFs, little else about the biology of noncanonical ORFs can be inferred from their lack of detection in MS data. We cannot conclude that proteins expressed from noncanonical ORFs are less stable than canonical proteins, that they are targeted for degradation at higher rates, or that they are less likely to be functional, except to the extent that low expression already justifies these inferences.

A majority of the yeast noncanonical translatome, and a small portion of the canonical, consist of evolutionarily young ORFs with little evolutionary conservation, classified as “evolutionary transient ORFs” in the Wacholder et al. dataset.^7^ No transient ORFs were detected in MS data, not even canonical transient ORFs that are well characterized. Evolutionary transient ORFs are both abundant in the genome and biologically significant, with some playing important roles in conserved pathways despite their short evolutionary lifespans.^7^ Though we were unable to detect them in MS data, numerous proteins expressed from evolutionarily transient ORFs are found to be present in the cell in microscopy studies.^7^ The biology of the vast majority of these ORFs are poorly understood; most have never been studied in any depth. Bottom-up MS, using currently available approaches, does not appear useful for identifying the evolutionarily transient ORFs most likely to have interesting biological roles.

There is considerable variability across studies that attempt to detect noncanonical proteins using MS, with some reporting detection of hundreds of proteins while others, as in this study, find many fewer.^10,13,15,18,21,31,36,60–62^ This could partly reflect biological differences between the cell types and species analyzed. However, there is also great variation in statistical approach. For example, though it is recommended for studies of noncanonical proteins to estimate a class-specific FDR among the noncanonical proteins themselves^24,63^, some studies control confidence using a whole-proteome FDR (including both canonical and noncanonical). Setting a strict whole-proteome FDR does not guarantee a low FDR among inferred noncanonical detections. In this study we found that, had we set a 1% whole-proteome FDR rather than controlling FDR among noncanonical proteins specifically, we would have produced a list of noncanonical protein detections comprised mostly of apparent false positives. This approach is made worse, moreover, when multiple datasets are analyzed independently, each using a 1% threshold, and then all hits are reported in a combined list. True detections are more likely to be shared between datasets than false positives, so the merged list will have a greater fraction of false positives than any of the individual dataset lists.^26^ To the extent that these practices are common, the published literature may paint a misleading picture of the ease of detecting ribo-seq supported noncanonical proteins. We believe that these issues can be addressed largely by following existing guidelines for FDR-based analyses and constructing adequate unbiased decoy sets. For example, the Human Proteome Project guidelines 3.0 state that, if multiple datasets are analyzed in a study, an FDR should be calculated on the combined dataset.^64^ Directly comparing the distribution of confidence scores among predicted noncanonical proteins and their unbiased decoys among all datasets provides a clear picture of the extent to which noncanonical proteins can be genuinely detected.

How, then, can we use shotgun mass spectrometry experiments to help us understand the biology of translated noncanonical ORFs and their potential protein products? We draw several lessons that may be applicable beyond yeast. For small-scale discovery of new protein-coding genes, the shotgun MS approach still provides value. Most noncanonical detections identified in this study were found on the basis of just one or two PSMs; additional support that these were genuine detections was provided from translation and evolution data. This suggests that further MS experiments conducted in a wide range of conditions will likely yield new discoveries of proteins that can be detected only rarely. To maximize these rare discoveries, it will be helpful to analyze MS data using different parameters, considering in particular a wide variety of post translational modifications. Given the short length of most noncanonical proteins, and that some noncanonical proteins lack tryptic peptides suitable for detection, it will likely also be helpful to use multi-enzyme digests or alternatives to digestion to maximize the probability that each noncanonical protein has at least one detectable peptide.^44^ However, we do not believe that these approaches alone will enable large-scale detection of noncanonical proteins such that shotgun MS will be useful for validating the presence (or absence) of most noncanonical proteins predicted by ribosome profiling experiments. As the vast majority of noncanonical proteins are outside the window of length and expression level in which canonical proteins are typically detected, technical advances that substantially improve sensitivity for small, low-abundance proteins may be needed for shotgun MS to serve this purpose.^22^ The three MS studies we examined here performed experimental enrichment of shorter and less abundant proteins, and further developments along these lines should facilitate broader noncanonical protein detection. We conclude that, while MS analysis of yeast ribo-seq supported noncanonical ORFs has some utility, it also has major limitations: it misses noncanonical proteins likely to be of biological interest, including an entire class of translated element, the evolutionarily transient ORFs. Targeted techniques for protein detection, such as microscopy^65^, Western blot, and top-down proteomics^60^, are more sensitive at detecting small proteins, but lack the convenience of untargeted bottom-up MS in being able to readily search for unannotated proteins predicted from an entire genome, transcriptome or translatome of a species. New technological developments in MS, and future innovations such as protein sequencing^66^, are needed to better assess the cellular presence and abundance of the great majority of proteins potentially encoded by the noncanonical translatome.

## Methods

### Mass spectrometry search

All mass spectrometry data files were taken from three studies. The He et al. 2018^29^ dataset PXD008586 and Gao et al. 2021 dataset PXD001928 were downloaded from PRIDE. The Sun et al. 2022^31^ dataset PXD028623 was downloaded from IPROX. These datasets were searched using all proteins predicted to be encoded from the full reference translatome described in Wacholder et al. 2023.^7^ The sequence database was supplemented with all canonical proteins not included in the Wacholder et al. 2023 dataset. Canonical proteins are those annotated as “verified”, “uncharacterized” or “transposable element” in the August 3, 2022 update of the Saccharomyces Genome Database annotation.^33^

Searches were conducted using the MSFragger program.^32^ Unless otherwise indicated, the following parameters were used: 20 ppm precursor mass tolerance, two enzymatic termini required, up to two missed cleavages allowed, clipping of the N-terminal methionine as a variable modification, methionine oxidation as a variable modification, cysteine carbamidomethylation as fixed modification, peptide digestion lengths from 7 to 50 amino acids, peptide masses from 350 to 1800 Daltons, a maximum fragment charge of 2. For the He et al. 2018 dataset and the Gao et al. 2021 dataset, fragment mass tolerance was set at 1 Da, while for the Sun et al. 2022 dataset fragment mass tolerance was set at 20 ppm; these settings reflect the instruments and settings used and were found to give the most canonical protein detections. Most experiments used trypsin as the digestive enzyme, but some of the experiments in Gao et al. 2021 were conducted using LysC; these experiments were analyzed using a separate parameter file setting LysC as the enzyme. After running MSFragger on each spectra file from the three studies, all output files, consisting of lists of PSMs and their properties, were concatenated together; analyses were done on PSMs pooled from all experiments.

Unless otherwise specified, FDR was calculated in a class-specific manner (i.e., specific to canonical or noncanonical ORFs) by dividing the number of decoy proteins within the class that were detected at the expect score threshold from the number of target proteins in the class detected at the threshold. A protein was considered detected at a given expect score threshold if had at least one unique PSM with an expect score below the threshold. Decoys were either default (reverse of protein database sequence) or reversed after the starting methionine, as indicated. Peptides were excluded if they belonged to more than one predicted protein. Peptides were also excluded from supporting noncanonical proteins if the exact peptide sequence existed in a canonical protein, regardless of whether it would be a tryptic peptide of that protein. Peptide-spectrum matches were excluded if the MSFragger hyperscore was less than 3 above the score for the next best peptide, in order to avoid using peptide-spectrum matches that did not uniquely support a single protein.

In two analyses, searches were instead conducted either using the MS-GF+ program^40^ or MaxQuant.^42^ All available parameters were set to be the same as in the MSFragger search, and decoys were reversed after the starting methionine. MS^2^Rescore^41^ was then run on MS-GF+ output files to rescore the results.

### Ribo-seq data

All ribo-seq data was taken from the analysis in Wacholder et al. 2023.^7^ This data included ribo-seq reads aggregated over 42 published studies and mapped to the *S. cerevisiae* genome. All reads are mapped to ribosome P-sites as described in Wacholder et al. 2023. A read was considered to map to an ORF only if the inferred P-site mapped to the first position of a codon in the reading frame of the ORF. The total read count for an ORF is the sum of reads mapping over all first codon positions, and the translation rate is the read count divided by the number of codons in the ORF.

### Homology analyses

BLAST analyses were conducted with default settings and a 10^-4^ e-value threshold to consider a match a homolog. BLAST searches conducted on NCBI databases were done on the NCBI website. Searches of the yeast genomes collected in Shen et al.^34^ were conducted using the BLAST command line tool on the genomes and annotations taken from that study.^68^ TBLASTN searches of *Saccharomyces* species genomes were conducted on genomes acquired from the following sources: *S. paradoxus* from Liti et al. 2009^69^, *S. arboricolus* from Liti et al. 2013^70^ (GCF_000292725.1), *S. jurei* from Naseeb et al. 2018^71^ (GCA_900290405.1), and *S. mikatae*, *S. uvarum*, *S. eubayanus* and *S. kudriavzevii* from Scannell et al. 2011.^72^ These genome were also used to make sequence alignments. All sequence alignments were generated using the MAFFT tool on the European Bioinformatics Institute website.^73^

### Peptide Analysis

To analyze the factors predicting MS detectability at the peptide level, a list of all possible tryptic peptides was constructed. For each ORF in the protein database, a set of possible peptides was constructed following the same rules as used for the MSFragger analysis: two enzymatic termini (or protein ends) were required, up to two missed cleavages were allowed, clipping to the N-terminal methionine was a variable modification, and methionine oxidation was a variable modification. As in the MSFragger analysis, peptides were restricted to 7 to 50 aa and peptide masses from 350 to 1800 Daltons. Out of this list of theoretical peptides, the peptides that were detected in the MS analysis at a 10^-6^ expect score threshold in at least one experiment were identified.

We used simulations to estimate the proportion of noncanonical proteins that would be detected if noncanonical peptides had the same detection probability as noncanonical proteins. Each simulation was conducted by randomly selecting noncanonical peptides at the canonical peptide detection rate and counting the proportion of noncanonical proteins with at least one selected peptide. The reported proportion is the average over ten simulations.

### Analyzing effect of peptide sequence and transmembrane domains

Every tryptic peptide was assessed for estimated detectability using the AP3 algorithm.^53^ Settings for peptide digestion were matched to that of the MS analysis and the pre-trained *S. cerevisiae* model provided with the program was used for scoring. Peptides were binned by AP3 detectability score, with ten intervals evenly spaced between 0 and 1, to construct figures 7C and 7D. We also used these intervals to estimate the proportion of canonical peptides that would be detected if canonical peptides had the same distribution of detectability scores as noncanonical peptides. For each bin, the canonical peptide detection probability was estimated as the rate of canonical peptide detection (MSFragger expect score < 10^-6^) within the bin. We then took the expected value of canonical peptide detections if the frequency distribution of canonical peptides among bins matched that of noncanonical peptides, and divided this count by the number of canonical peptides actually detected.

Every protein was assessed for transmembrane domains using TMHMM.^54^ For each protein, the proportion of amino acids assigned by TMHMM to a transmembrane domain was calculated if at least one transmembrane helix was predicted; otherwise the proportion was set at zero. To construct Figure 7D, proteins were binned by transmembrane proportion, with each bin covering an interval of 0.1, and each peptide was assigned to the same bin as its associated protein. The difference in distribution between canonical proteins and noncanonical proteins larger than 50 aa was assessed using a chi-squared test on a contingency table containing the counts of each class in each bin.

## Data availability

All data and code used in this study can be found on Figshare (https://doi.org/10.6084/m9.figshare.24026367).

## Acknowledgments

We thank Jiwon Lee for helpful feedback. This work was supported by funds provided by the Searle Scholars Program to A.-R.C. and the National Institute of General Medical Sciences of the National Institutes of Health grant DP2GM137422 (awarded to A.-R.C.).

## Author contributions

Conceptualization, A.W. and A.-R.C. Methodology, A.W., A.-R.C. Investigation, A.W. Writing – Original Draft, A.W. Writing – Review & Editing, A.W., A.-R.C. Supervision, A.-R.C.

## Supporting Information Legends

**Supplementary Figure 1: Spectra supporting detection of noncanonical protein YFR035W-A.** Two spectra with stronger MSFragger expect scores than all decoys are shown. At top, the full protein sequence is shown, with peptide coverage colored in red. Spectra were visualized using the Universal Spectrum Identifier (USI) tool on ProteomeExchange^67^ for the USIs given in Supplementary Table 1. The data underlying this Figure can be found in S1 Data.

**Supplementary Figure 2: Spectra supporting detection of noncanonical protein YMR106W-A.** A single PSM (top left) had a stronger MSFragger expect score than all decoys, while three others had stronger scores than the next strongest decoy. At top, the full protein sequence is shown, with peptide coverage colored in red. Spectra were visualized using the Universal Spectrum Identifier tool on ProteomeExchange for the USIs given in Supplementary Table 1. The data underlying this Figure can be found in S1 Data.

**Supplementary Figure 3: Spectrum supporting detection of noncanonical protein YIL059C.** A single PSM had a stronger MSFragger expect score than all decoys when a single non-enzymatic end was allowed. At top, the full protein sequence is shown, with peptide coverage colored in red. The spectrum was visualized using the Universal Spectrum Identifier tool on ProteomeExchange for the USI given in Supplementary Table 1. The data underlying this Figure can be found in S1 Data.

**Supplementary Figure 4: Claimed noncanonical detections in previous study have similar scores to decoys.** For each protein and decoy passing the detection threshold in the Sun et al.^31^ study, the strongest score among all PSMs associated with the protein or decoy is indicated. All scores and protein classifications were taken from output files of Sun et al.^31^ downloaded from IRPOX (PXD028623); we combined all PSMs from 11 different output files to create the plot. Each output file contains the PSMs passing a 1% FDR threshold, set at the whole-proteome level (i.e., not distinguishing canonical from noncanonical), in analyses conducted by Sun et al. using pFind. Each output file was individually thresholded at a 1% FDR in the Sun et al. analysis and noncanonical proteins passing this threshold in any file were inferred to be detected. Lower scores indicate higher confidence given by the MS algorithm. The observation that decoys and claimed noncanonical detections have similar scores suggests that many claimed noncanonical detections (indicated in blue) may be false positives. Merging multiple lists of inferred detections that were each individually generated at a 1% FDR is expected to result in a combined list with a much higher FDR^26^, which, together with the use of a proteome-wide rather than noncanonical-specific FDR, can help explain why many noncanonical proteins were inferred to be detected despite scoring similarly to decoys. The data underlying this Figure can be found in S1 Data.

**Supplementary Figure 5: Spectra supporting detection of noncanonical protein YNL155C-A.** Spectra for the four strongest matches for YNL155C-A. At top, the full protein sequence is shown, with peptide coverage colored in red. The spectra were visualized using the Universal Spectrum Identifier tool on ProteomeExchange for the USIs given in Supplementary Table 1. The data underlying this Figure can be found in S1 Data.

**Supplementary Figure 6: Very few noncanonical proteins have high-confidence PSMs.** The best peptide-spectrum match MSFragger expect score for each noncanonical protein and decoy in the database. Lower scores indicate stronger matches. The data underlying this Figure can be found in S1 Data.

**Supplementary Figure 7: Translation rate declines with ORF size.** A) Average log in-frame ribo-seq read count per codon among canonical ORFs of different size classes. B) Average log in-frame ribo-seq read count per codon among noncanonical ORFs of different size classes. The data underlying this Figure can be found in S1 Data.

**Supplementary Figure 8: Proteins with more transmembrane content are less detectable by MS.** The proportion of peptides detected, among all eligible for detection, for canonical proteins binned by proportion of amino acids in predicted transmembrane domains. Predictions were made using TMHMM.^54^ The first bin includes only proteins with no transmembrane domain predicted. The data underlying this Figure can be found in S1 Data.

**Supplementary Figure 9: Evolutionarily transient canonical proteins found at similar rates to decoys.** Predicted proteins and decoys detected in MS data at a range of expect-score thresholds, among canonical proteins identified as evolutionarily transient in Wacholder et al. 2023^7^, using the standard MSFragger approach. The data underlying this Figure can be found in S1 Data.

**S1 Data: Data underlying figures 1, 2, 3, 4, 5, 6, 7, and S1, S2, S3, S4, S5, S6, S7, S8, S9.**

